# Aicardi-Goutières Syndrome gene *Rnaseh2c* is a metastasis susceptibility gene in breast cancer

**DOI:** 10.1101/550418

**Authors:** Sarah K. Deasy, Ryo Uehara, Suman K. Vodnala, Howard H. Yang, Randall A. Dass, Ying Hu, Maxwell P. Lee, Robert J. Crouch, Kent W. Hunter

**Author notes:** Corresponding Author: Kent W. Hunter.

## Abstract

Breast cancer is the second leading cause of cancer-related deaths in the United States, with the majority of these deaths due to metastatic lesions rather than the primary tumor. Thus, a better understanding of the etiology of metastatic disease is crucial for improving survival. Using a haplotype mapping strategy in mouse and shRNA-mediated gene knockdown, we identified *Rnaseh2c*, a scaffolding protein of the heterotrimeric RNase H2 endoribonuclease complex, as a novel metastasis susceptibility factor. We found that the role of *Rnaseh2c* in metastatic disease is independent of RNase H2 enzymatic activity, and immunophenotyping and RNA-sequencing analysis revealed engagement of the T cell-mediated adaptive immune response. Furthermore, the cGAS-Sting pathway was not activated in the metastatic cancer cells used in this study, suggesting that the mechanism of immune response in breast cancer is different from the mechanism proposed for Aicardi-Goutières Syndrome, a rare interferonopathy caused by RNase H2 mutation. These results suggest an important novel, non-enzymatic role for RNASEH2C during breast cancer progression and add *Rnaseh2c* to a panel of genes we have identified that together could determine patients with high risk for metastasis. These results also highlight a potential new target for combination with immunotherapies and may contribute to a better understanding of the etiology of Aicardi-Goutières Syndrome autoimmunity.

**Author Summary:** The majority of breast cancer-associated deaths are due to metastatic disease, the process where cancerous cells leave the primary tumor in the breast and spread to a new location in the body. To better understand the etiology of this process, we investigate the effects of gene expression changes in the primary tumor. In this study, we found that changing the expression of the gene *Rnaseh2c* changed the number of metastases that developed in the lungs of tumor-bearing mice. By investigating the enzyme complex *Rnaseh2c* is part of, RNase H2, we determined that *Rnaseh2c*’s effects may be independent of RNase H2 enzyme activity. Because *Rnaseh2c* is known to cause the autoimmune disease Aicardi-Goutières Syndrome (AGS), we tested whether the immune system is involved in the metastatic effect. Indeed, we found that the cytotoxic T cell response is important for mediating the effect that *Rnaseh2c* has on metastasis. Together these data indicate that *Rnaseh2c* expression contributes to a patient’s susceptibility to developing breast cancer metastasis and demonstrate that the immune system is involved in this outcome. The implications of this study suggest immunotherapy could be a viable treatment for breast cancer metastasis and may help inform the understanding of AGS and RNase H2 in cancer.

## Introduction

Breast cancer is the most frequently diagnosed female malignancy in the United States and the second leading cause of cancer-related female deaths [1]. Since the primary tumor is usually resected upon detection, most of this mortality is due to metastases. The 5-year survival for patients diagnosed with localized breast cancer is approximately 99%, indicating that detection and treatment is highly effective when the cancer remains in the breast tissue. However, 5-year survival for patients with overt distant metastases plummets to a dismal 27% [1]. These statistics indicate that the establishment of effective therapies that target and prevent metastasis is of critical importance. Although recent decades have seen significant advances in the development of methods for detection and treatment of primary breast cancer, these treatments are generally ineffective at dampening metastatic progression. Therefore, a better understanding of the mechanisms that regulate and drive metastasis is critical to improving patient outcome.

As part of the effort to understand the biological etiology of metastatic disease, we investigate the impact of genetic polymorphism on metastatic progression and susceptibility. Using a strategy that integrates population genetics analysis with a variety of genomic tools, we previously reported and characterized polymorphic genes that drive metastatic susceptibility. Since the majority of disease-associated polymorphisms are non-coding and are thought to influence gene expression rather than protein structure [2], we hypothesized that altering the expression of these key genes can affect metastatic outcome. This approach has generated a growing list of metastasis susceptibility and tumor progression genes with tumor-autonomous and/or stromal effects [3–11], including some with the potential to be actionable clinical targets.

Here, we report the identification of a ~ 1 megabase haplotype containing 74 genes on mouse chromosome 19 that associated with changes in pulmonary metastasis. Within this haplotype, five genes exhibited gene expression patterns significantly associated with metastasis, of which only one, *Rnaseh2c* (Entrez GeneID: 68209), exhibited a positive association with metastasis, therefore potentially being amenable to clinical inhibition. This gene encodes a scaffolding subunit of the Ribonuclease H2 (RNase H2) enzyme. RNase H2 initiates the removal of ribonucleotides that are incorporated into DNA, which can occur as single ribonucleotides or as longer stretches of RNA/DNA hybrids [12]. The RNase H2 complex is composed of a catalytic A subunit (encoded by *Rnaseh2a*, 69724) and a second scaffolding B subunit (*Rnaseh2b*, 67153) in addition to subunit C. In humans, mutations in any of these three genes result in an autosomal recessive neurological autoinflammatory disorder called Aicardi-Goutières Syndrome (AGS) [13]. This syndrome clinically mimics the phenotype of *in utero* viral infection and is characterized by basal ganglia calcification, progressive microcephaly with accompanying mental and motor retardation, and lymphocytosis and elevated Interferon α in the cerebral spinal fluid and serum at early points in disease progression which activates a type I Interferon signature [14–16]. In many cases, these children ultimately die early in life. RNase H2 mutations have also been identified in patients with another autoimmune disease, Systemic Lupus Erythematosus (SLE), though it is not known if these mutations are causative [17]. Given its important role in maintaining genomic integrity, a connection between RNase H2 and cancer is beginning to be investigated, with multiple studies having identified associations between expression of RNase H2 genes and various cancer types [18–25].

We report here the validation of *Rnaseh2c* as a bona fide metastasis susceptibility gene and reveal its novel function in promoting metastasis through interaction with the immune system, although through an alternate mechanism than that proposed for AGS. This knowledge may contribute to a better understanding of the biology underlying AGS and highlights a potential new target for combination with immune modulatory therapies to combat metastatic breast cancer. This data also adds to a growing panel of genes we have identified that together could help determine patients who are at high risk for developing metastasis.

## Results

### Identification of *Rnaseh2c* as a candidate metastasis susceptibility gene

Genetic mapping studies were performed to identify metastasis susceptibility gene candidates. A population of genetically diverse tumor-bearing mice was generated by crossing the FVB/NJ-TgN(MMTV-PyMT)^634Mul^ (hereafter, MMTV-PyMT) mouse model of metastatic breast cancer [26] to animals from the Diversity Outbred mouse mapping panel [27, 28]. 50 DO Generation 5 mice were crossed with MMTV-PyMT, as described in [29] (Fig 1A). Offspring from the DO x MMTV-PyMT crosses (n = 115) were aged to humane endpoint at which time primary tumors were weighed and surface pulmonary metastases counted. Each of the tumor-bearing mice were genotyped and the unique combinations of strain-specific haplotypes in the offspring genomes were calculated for each mouse. The haplotype analysis was integrated with the phenotype data to identify metastasis-associated regions. A total of 17 haplotypes on 11 chromosomes were found to be significantly associated with metastatic disease (FDR < 0.05) (S1 Fig A). Interestingly, two of the 17 haplotypes overlapped with regions previously identified in genetic studies of metastasis susceptibility: a 1.1 megabase haplotype on chromosome 9 that overlapped the peak of a metastasis susceptibility locus previously found to contain the metastasis susceptibility gene *Pvrl1* [7, 11]; and a 1.3 megabase haplotype on mouse chromosome 19 that overlapped with a previously identified metastasis susceptibility locus which includes the metastasis susceptibility gene *Sipa1* [30]. Due to the high density of genes in the chromosome 19 haplotype and the fact that modifier loci can contain multiple causative genes [31], we selected this region for further screening to identify additional metastasis susceptibility genes that might exist within this interval. Given that polymorphism effects are frequently on gene expression [2], primary tumor gene expression analysis was performed to identify genes whose expression correlated with pulmonary metastasis. This analysis identified five genes located in the region of interest: *Frmd8, Mus81, Rnaseh2c, Pola2*, and *Dpp3* (S1 Fig B). *Sipa1* was not identified by this screen, consistent with previous studies that demonstrated that the effect of *Sipa1* on metastasis was due to a missense polymorphism rather than an expression polymorphism [9]. Interestingly, analysis of patient tumor expression using the GOBO database [32] revealed that the genes within this signature, weighted for expression direction and relative contribution to the signature using the mouse-calculated hazard ratios and grouped by their cumulative signature score, are also able to predict distant metastasis-free survival in breast cancer patients of the Her2-enriched subtype (S1 Fig C), suggesting that our approach is able to identify haplotypes relevant to outcome in patients as well as in mice. *Rnaseh2c* was selected for further investigation due to it being the only candidate gene within the haplotype that was positively associated with metastasis, thus harboring the possibility that therapeutic inhibition could reduce metastatic burden.

**Fig 1.**
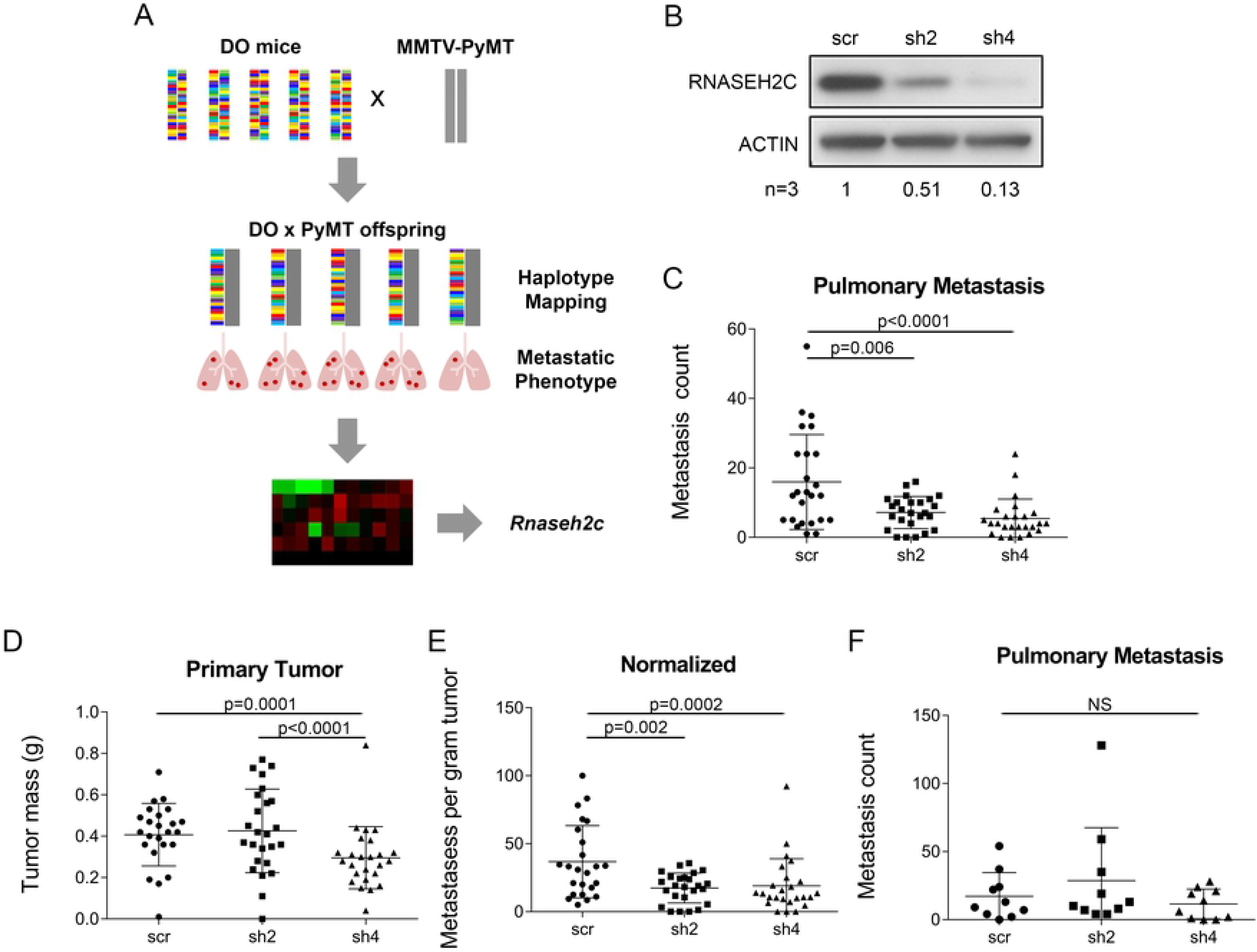
Knockdown of *Rnaseh2c* reduces pulmonary metastasis. (A) Mice from the DO Generation 5 were bred to the MMTV-PyMT transgenic model to induce tumorigenesis and metastasis in the offspring, which were used to identify strain-specific haplotypes associated with metastasis. A tumor expression microarray was used to identify genes within the identified haplotypes exhibiting changes in expression associated with changes in metastasis. *Rnaseh2c* was one of the identified candidate genes. (B) RNASEH2C protein expression by western blot upon shRNA-mediated knockdown. One representative blot is shown, average relative densitometry for three independent experiments is reported below. (C-E) Pulmonary metastases (C) and tumor mass (D) from orthotopically injected scramble control (scr) and knockdown cells were quantified at euthanasia and normalized (metastases per gram of tumor, E); average ± standard deviation, n = 25 mice per group in three independent experiments. (F) Tail vein experimental metastasis experiment using scramble and knockdown cells. Pulmonary metastases were counted at euthanasia; average ± standard deviation, n = 10 mice per group. NS – not significant

### Modulating expression of *Rnaseh2c* alters spontaneous metastasis

To investigate the role of *Rnaseh2c* as a metastasis modifier gene, shRNA-mediated knockdowns were performed in the mouse metastatic mammary tumor cell line Mvt1. Knockdown was confirmed by qRT-PCR (S2 Fig A), and the sh2 and sh4 cell lines were selected for future experiments based on western blot analysis (Fig 1B). To determine the effects on tumor growth and progression, knockdown Mvt1 cells were orthotopically injected into the mammary fat pad of syngeneic FVB mice. Both knockdown cell lines exhibited significantly reduced numbers of spontaneous pulmonary metastases (Fig 1C), indicating that changes in *Rnaseh2c* expression affect the cell’s ability to metastasize. In addition, the sh4 cell line showed reduced primary tumor mass (Fig 1D), however both knockdown cell lines maintained the significant reduction in metastasis counts when normalized to their matched primary tumors (Fig 1E). To verify *Rnaseh2c*’s role as a bona fide metastasis modifier, we knocked down *Rnaseh2c* in a second mouse metastatic mammary cancer cell line, 4T1 (S3 Fig A&B). In contrast to the primary tumor effect in the Mvt1 cell line, sh2 reduced while sh4 increased primary tumor mass of the 4T1 cells (S3 Fig C), suggesting that the effect of the shRNAs on tumor mass was independent of *Rnaseh2c* expression. Again, both knockdown cell lines showed reductions in pulmonary metastases that were maintained upon normalization (S3 Fig D&E), demonstrating that the effect of *Rnaseh2c* on metastasis is not cell line-specific.

Since reduction in metastatic burden can result from effects along many points of the metastatic cascade, an experimental (tail vein) metastasis assay was performed to determine whether reduced *Rnaseh2c* expression affects the final extravasation and colonization steps. No significant difference in the number of pulmonary metastases was observed between knockdown and control conditions (Fig 1F). This indicated that reduction of *Rnaseh2c* expression in tumor cells did not affect metastatic colonization after a bolus injection into the venous circulation.

To complement the knockdown assays, we also investigated the effect of increased *Rnaseh2c* expression on metastasis. Total protein expression (combined endogenous and FLAG-tagged exogenous protein) was increased approximately threefold in Mvt1 cells that were transduced with an *Rnaseh2c* expression vector compared with cells transduced with an empty vector (Fig 2A), in agreement with the qRT-PCR results (S2 Fig B). Interestingly, endogenous RNASEH2C protein levels appeared to be lower in the presence of exogenous expression, suggesting that there may be mechanisms to prevent excessive RNASEH2C protein in cells. In agreement with the results from the knockdown metastasis assays, increased expression resulted in more pulmonary metastases without affecting primary tumor mass (Fig 2 B-D). Together these data confirm *Rnaseh2c* as a metastasis modifier gene, specifically as a metastasis promoter.

**Fig 2.**
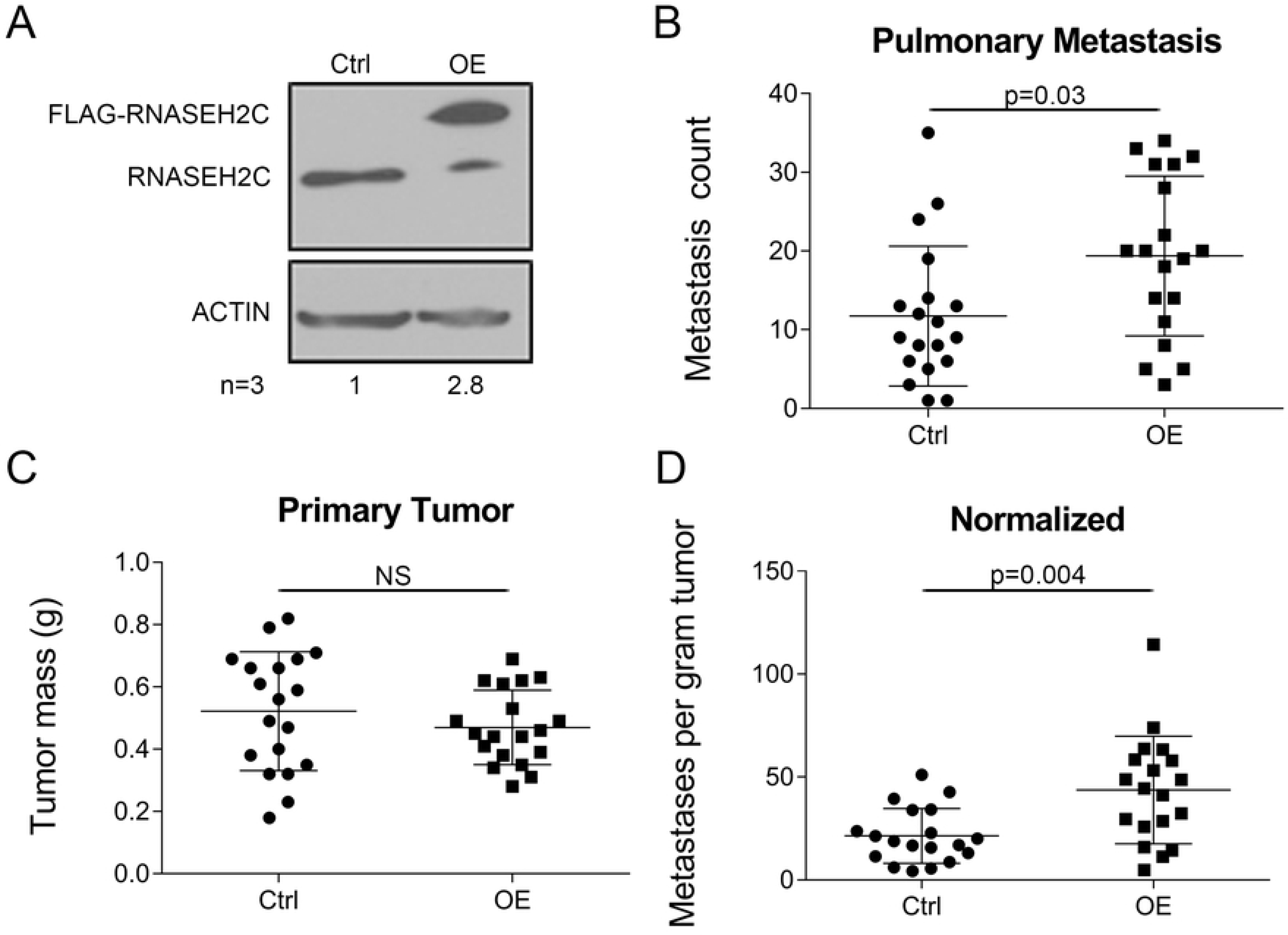
Overexpression of *Rnaseh2c* increases pulmonary metastasis. (A) RNASEH2C protein expression by western blot following exogenous overexpression (OE). One representative blot is shown, average relative densitometry for three independent experiments is reported below; OE lane is combined endogenous and exogenous tagged RNASEH2C expression. (B-D) OE cell line was orthotopically injected along with empty vector control (Ctrl) as described. Pulmonary metastases (B) and tumor mass (C) were quantified at euthanasia and normalized (metastases per gram of tumor, D); average ± standard deviation, n = 19 mice per group in two independent experiments. NS – not significant

### Reduced RER enzyme activity upon *Rnaseh2c* knockdown is not the driver of the metastatic effect

RNase H2 is a heterotrimeric complex composed of a catalytic subunit, RNASEH2A, and two scaffolding subunits, RNASEH2B and RNASEH2C. All three subunits for required for proper enzyme function [33, 34]. We therefore tested whether knockdown of *Rnaseh2c* reduced cellular enzyme activity. An *in vitro* RNase H2 enzyme activity assay was performed by incubating cell lysates from scramble and sh4 knockdown cell lines with an exogenous labelled DNA 12-mer containing a single ribonucleotide to test the RNase H2-specific ribonucleotide excision repair (RER) activity. *Rnaseh2c* knockdown resulted in approximately 30% substrate cleavage compared to scramble (Fig 3A), indicating that limiting the available RNASEH2C protein reduces RNase H2 RER enzyme activity in Mvt1 cells.

**Fig 3.**
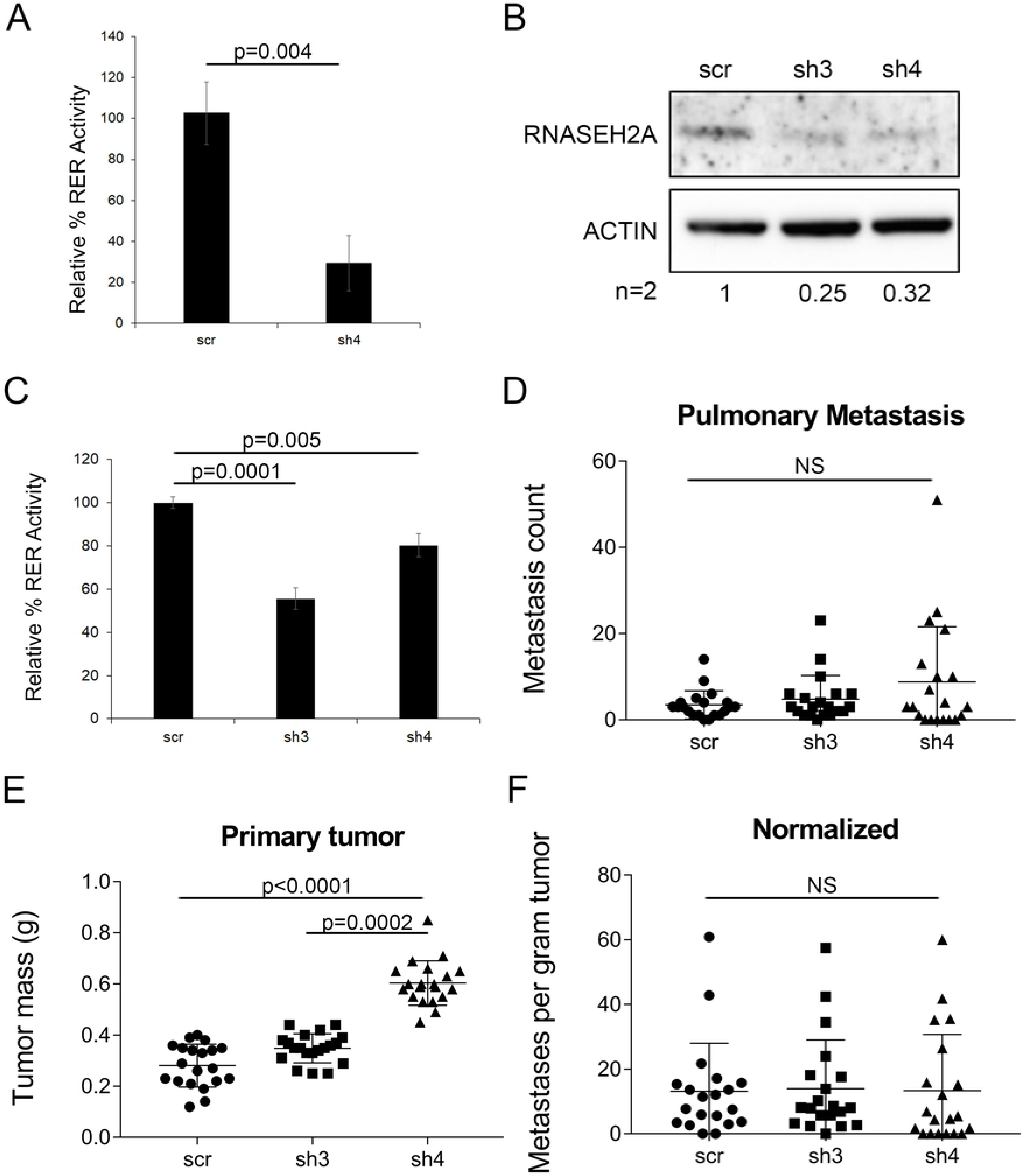
Knockdown of *Rnaseh2a* does not phenocopy the metastatic effect of *Rnaseh2c*. (A) Percent ribonucleotide excision repair (RER) activity was measured in Mvt1 scr and sh4 *Rnaseh2c* knockdown cells. Average ± standard deviation of three experiments. (B) RNASEH2A protein expression by western blot. One representative blot is shown, average relative densitometry for two independent experiments is reported below. (C) Percent ribonucleotide excision repair (RER) activity was measured in Mvt1 scr and *Rnaseh2a* knockdown cells. Average ± standard deviation of three experiments. (D-F) Control and *Rnaseh2a* knockdown cells were used in spontaneous metastasis assays. Pulmonary metastases (D) and tumor mass (E) were quantified at euthanasia and normalized (metastases per gram of tumor, F); average ± standard deviation, n = 20 mice per group in two independent experiments. NS – not significant

To test if this reduction in enzyme activity is the reason for reduced metastasis, we evaluated the effects of knocking down the catalytic subunit, *Rnaseh2a*, in Mvt1 cells. Knockdown was evaluated by qRT-PCR (S2 Fig C) and western blot (Fig 3B). RER activity analysis revealed that the *Rnaseh2a* knockdown significantly reduced enzyme RER activity (Fig 3C). Spontaneous metastasis assays with the *Rnaseh2a* knockdown cells produced an increase in primary tumor mass, however there was no significant difference in either the metastasis count or upon normalization (Fig 3 D-F). Thus, the effect of *Rnaseh2c* knockdown on metastasis is likely not the result of the reduced RER activity and may suggest an enzyme-independent function for the subunit in metastasis.

### *Rnaseh2c* knockdown does not affect Mvt1 proliferation, apoptosis, cell cycle progression, or activate a DNA damage response

Given that reduction in enzyme RER activity did not consistently correlate with reduced metastasis, we tested whether *Rnaseh2c* knockdown cells exhibit downstream effects of RNase H2 dysfunction. Cellular effects reported in the literature to be altered upon RNase H2 enzyme disruption were tested in *Rnaseh2c* knockdown cell lines and tumors. First, previous reports indicate that dysfunctional RNase H2 results in reduced cell proliferation both in mouse fibroblasts [35] and in fibroblasts from AGS and SLE patients [17]. Therefore, proliferation upon *Rnaseh2c* knockdown was evaluated both in cell lines (S4 Fig A) and in tumors (Fig 4A, S4 Fig C). A trending reduction in Ki67 staining was observed *in vivo* for the knockdown tumors, however this trend was not significant. Apoptosis was also measured in cell lines (S Fig 4B) and in tumors (Fig 4A, S4 Fig C) by cleavage of caspase 3; no differences were observed. Together these results suggest that the level of RNase H2 disruption achieved by the *Rnaseh2c* knockdown does not extend to affecting cell proliferation and apoptosis in this cell line. The additional *in vivo* evidence further confirms that the effect of *Rnaseh2c* expression on metastasis is not primarily through altering tumor growth.

**Fig 4.**
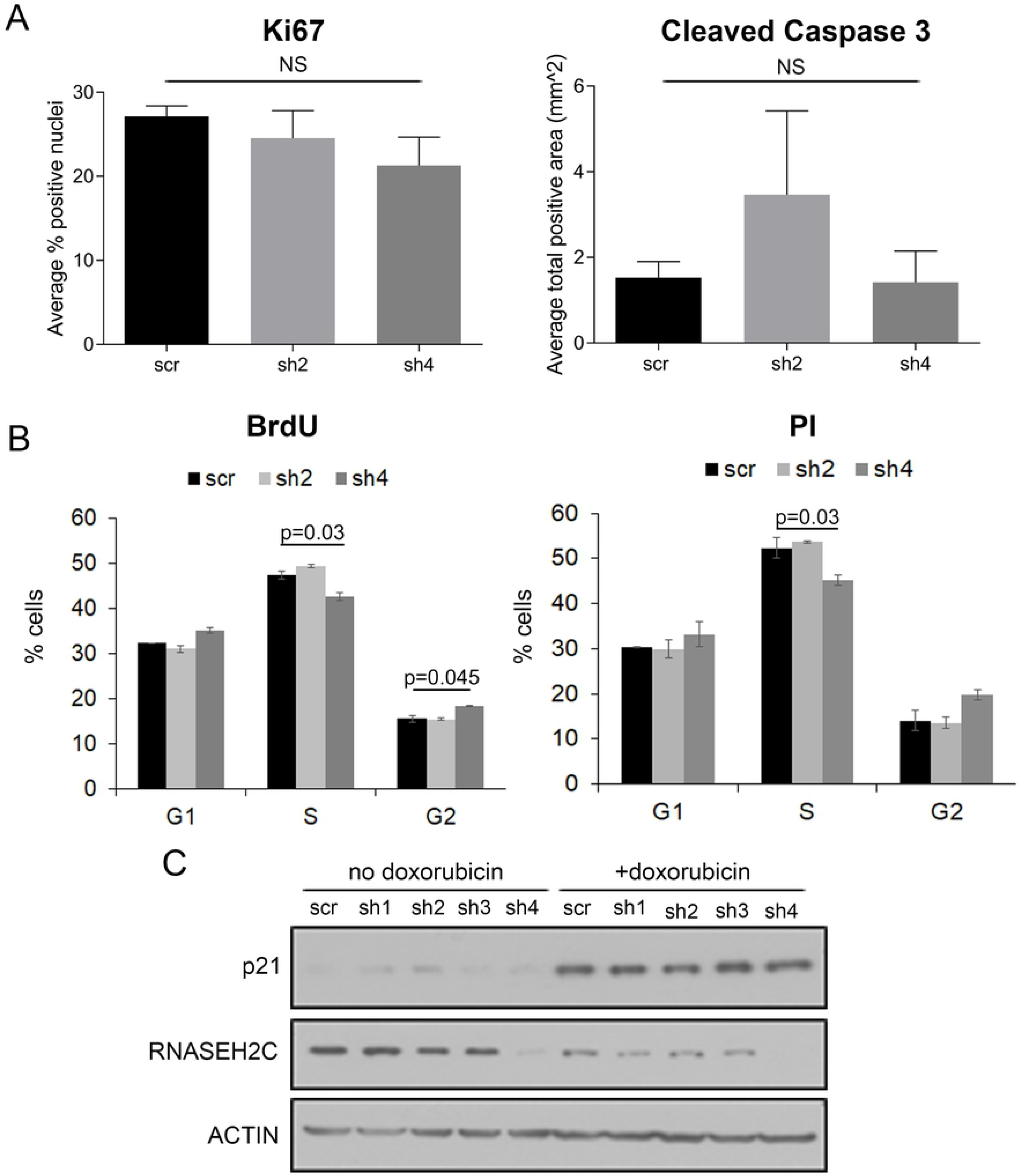
*Rnaseh2c* knockdown does not affect Mvt1 proliferation, apoptosis, cell cycle progression, or induce a DNA damage response. (A) Quantification of Ki67 (left) and Cleaved Caspase 3 (right) staining by IHC of tumor sections as the average ± standard deviation of one section from each of three independent tumors. (B) Unsynchronized cells were labeled with BrdU (left) and stained with propidium iodide (PI, right) to analyze cell cycle progression. Average ± standard deviation of duplicate samples. (C) Cells ± doxorubicin treatment (1μM for 24 hr) were analyzed for p21 protein expression in response to DNA damage. NS – not significant

Cell lines were also assayed for differences in cell cycle progression, which is reportedly stalled upon RNase H2 dysfunction in mouse and human fibroblasts [17, 35]. Both propidium iodide staining and BrdU incorporation indicated only minimal accumulation of *Rnaseh2c* sh4 knockdown cells in the G2 phase with no other significant differences observed (Fig 4B). The cell cycle disruption observed in mouse-derived fibroblasts is reportedly accompanied by a concomitant increase in p53-inducible genes, notably p21, indicating activation of the DNA damage response pathway [35]. Expression of p21 was not found to be upregulated upon *Rnaseh2c* knockdown (Fig 4C), though expression could be induced by treatment with doxorubicin, a chemotherapeutic that causes DNA damage partially by inducing replication stress through inhibiting topoisomerase II [36]. Given this mechanism of action, we hypothesized that our knockdown cells may be more sensitive to doxorubicin treatment due to cells already experiencing replication stress because of a genomic ribonucleotide burden resulting from disrupted RNase H2 activity. Therefore, we treated control and knockdown cells with increasing concentrations of doxorubicin and measured cell viability. However, the cell lines were equally sensitive to the doxorubicin treatment (S4 Fig D). In agreement with the p21 data, we also did not observe an induction of γ-H2AX signal in our knockdown cells (S5 Fig), suggesting the knockdown does not elicit a DNA damage response.

### Expression of RNASEH1 is upregulated in response to *Rnaseh2c* knockdown

Since RNase H2 resolves RNA/DNA hybrids in the cell, we stained *Rnaseh2c* knockdown cells with the S9.6 antibody which recognizes these nucleic acid structures to determine how they change in response to knockdown. In agreement with previous reports [37], the majority of staining in scramble control cells was observed in the cytoplasm with only a small amount concentrated in regions of the nucleus (S6 Fig A). Unexpectedly, in knockdown cells, staining was slightly reduced in the sh2 and drastically reduced in the sh4 cells including in the nuclei (S6 Fig A), suggesting that hybrids were not accumulating despite reduced RNase H2 enzyme activity. We hypothesized that this could be due to a compensatory upregulation of *Rnaseh1*, which also removes longer RNA moieties from RNA/DNA hybrids. We found that RNASEH1 expression was indeed increased upon *Rnaseh2c* knockdown (S6 Fig B). To test whether *Rnaseh2c* knockdown alters enzyme activity against RNA/DNA hybrids, we repeated the enzyme activity assay using an oligo substrate composed of an RNA strand hybridized to a DNA strand. This analysis revealed no difference between the control and *Rnaseh2c* knockdown cells (S6 Fig C), suggesting that RNase H1 upregulation helps maintain activity against RNA/DNA hybrids.

Together, these data show that despite a significant reduction in RNase H2 enzyme activity, the expected effects of enzyme dysfunction are not observed in our cancer cells, suggesting there may be other enzyme-independent effects taking place. We therefore expanded our investigation into other potential effects of reduced enzyme activity.

### Gene expression profiling reveals alterations to T cell immune response pathways upon *Rnaseh2c* knockdown

Given that the expected effects of enzyme dysfunction were not observed upon *Rnaseh2c* knockdown, we next profiled the gene expression landscape in response to *Rnaseh2c* knockdown to identify global transcriptional effects. We performed mRNA sequencing of the scramble control and sh4 knockdown tumors which enabled us to capture gene expression changes from both tumor and stroma in the *in vivo* context. GSEA analysis revealed that genes of the Inflammatory Response gene set approached significant enrichment (nominal p-value = 0.07) in the sh4 knockdown tumors compared to scramble, with a significant enrichment of the Interferon Gamma Response and Interferon Alpha Response gene sets (Fig 5A). For more specific pathway annotations, genes with expression changes of at least two-fold in either direction (249 genes, 169 of which exhibited a positive fold change) were assigned to biological pathways and analyzed by Ingenuity Pathway Analysis (S1 File). Of the top 25 significantly altered pathways, 17 were related to T cell responses (Table 1). Furthermore, more than half (8 of 17) of these pathways produced positive Z-scores, indicating that the direction of transcript change is consistent with the pathway being activated in knockdown tumors.

**Fig 5.**
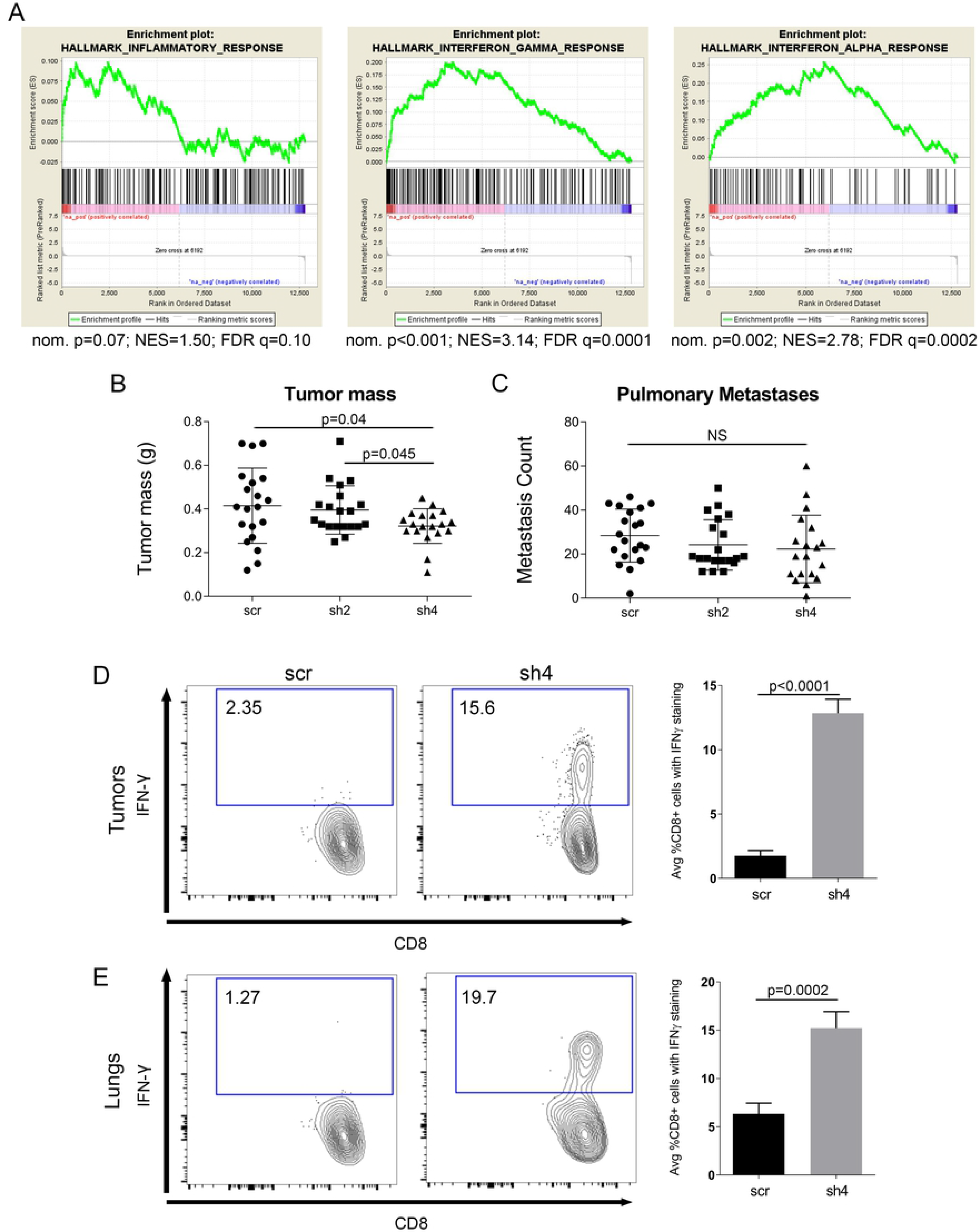
T cells mediate the effect of *Rnaseh2c* knockdown on metastasis. (A) GSEA analysis of RNA-sequencing data comparing sh4 with scr tumors in hallmark pathways related to inflammation and interferon signaling. nom. p: nominal p-value; NES: normalized enrichment score; FDR q: false discovery rate q-value. (B) Tumor mass in athymic nude mice that received Mvt1 cells with a knockdown of *Rnaseh2c*; average ± standard deviation. (C) Metastasis counts for the mice from (B); average ± standard deviation, n = 20 mice per group in two independent experiments. (D) Proportion of Interferon γ (IFN-γ)-producing CD8+ T cell that were isolated from tumors of FVB mice that received orthotopic injection with either scr or sh4 cells; average ± SEM. (E) Proportion of Interferon γ (IFN-γ)-producing CD8+ T cell that were isolated from metastatic lungs of the mice shown in (D); average ± SEM. n=5 mice per group with samples from each mouse analyzed in triplicate. NS – not significant

**Table 1.** Top 25 altered pathways as determined by RNA-sequencing IPA analysis

| Ingenuity Canonical Pathways | p-value | ratio | z-score |
| --- | --- | --- | --- |
| CTLA4 Signaling in Cytotoxic T Lymphocytes | 2.45E-08 | 0.101 | -- |
| T Cell Receptor Signaling | 7.24E-06 | 0.073 | -- |
| iCOS-iCOSL Signaling in T Helper Cells | 1.17E-04 | 0.058 | 2.236 |
| CD28 Signaling in T Helper Cells | 1.86E-04 | 0.054 | 1 |
| Hematopoiesis from Pluripotent Stem Cells | 8.91E-04 | 0.085 | -- |
| Primary Immunodeficiency Signaling | 9.55E-04 | 0.083 | -- |
| PKCθ Signaling in T Lymphocytes | 1.26E-03 | 0.046 | 2 |
| Th1 Pathway | 1.41E-03 | 0.045 | 2.236 |
| Th1 and Th2 Activation Pathway | 1.48E-03 | 0.038 | -- |
| Agranulocyte Adhesion and Diapedesis | 1.78E-03 | 0.037 | -- |
| CDK5 Signaling | 2.14E-03 | 0.051 | -0.447 |
| Th2 Pathway | 2.40E-03 | 0.041 | 2.236 |
| Calcium-induced T Lymphocyte Apoptosis | 2.63E-03 | 0.064 | 2 |
| Cytotoxic T Lymphocyte-mediated Apoptosis of Target Cells | 3.09E-03 | 0.094 | -- |
| OX40 Signaling Pathway | 5.13E-03 | 0.053 | -- |
| Neuroprotective Role of THOP1 in Alzheimer's Disease | 5.75E-03 | 0.075 | -- |
| Regulation of IL-2 Expression in Activated and Anergic T Lymphocytes | 5.89E-03 | 0.051 | -- |
| Phospholipase C Signaling | 6.03E-03 | 0.030 | 1 |
| Role of NFAT in Regulation of the Immune Response | 6.76E-03 | 0.033 | 2.449 |
| Communication between Innate and Adaptive Immune Cells | 8.91E-03 | 0.045 | -- |
| Human Embryonic Stem Cell Pluripotency | 9.77E-03 | 0.035 | -- |
| Granzyme A Signaling | 1.29E-02 | 0.105 | -- |
| Nur77 Signaling in T Lymphocytes | 1.45E-02 | 0.054 | -- |
| Type I Diabetes Mellitus Signaling | 1.74E-02 | 0.037 | -- |
| Tight Junction Signaling | 1.86E-02 | 0.030 | -- |

*In silico* analysis using ImmQuant software also supported these findings. ImmQuant uses RNA-sequencing data and an immune deconvolution algorithm to predict the presence of immune cell populations in heterogeneous tissues based on comparison to reference gene expression profiles of defined inflammatory states in mouse [38]. This analysis predicted an increase in *Rnaseh2c* knockdown tumors of macrophages and dendritic cells, which may be important for facilitating the observed adaptive immune response. In addition, the largest predicted increase was for CD8+ T cells (S7 Fig). Together these data suggest that the immune system, and CD8+ T cells in particular, are important for the effect that *Rnaseh2c* has on limiting metastasis.

### A CD8+ T cell response is involved in reducing cellular metastatic capacity upon *Rnaseh2c* knockdown

Individuals with inherited RNase H2 mutations develop the neurological autoinflammatory disorder AGS [13]. We were therefore intrigued to find by RNA-sequencing that immune response-related genes were transcriptionally activated upon *Rnaseh2c* knockdown in breast cancer. To investigate whether this immune activation is important for metastatic outcome, orthotopic metastasis assays were performed using athymic nude mice, which lack mature T cells. While the previously observed significant reduction in sh4 primary tumor mass was maintained (Fig 5B), the pulmonary metastasis effect was ablated (Fig 5C) indicating that the *Rnaseh2c* metastasis effect functions through engagement of T cell-mediated immunity.

To further delineate which T cell populations were involved, immunophenotyping was performed using tissue from the primary tumors, metastatic lungs, and spleens of mice orthotopically injected with the scramble control or knockdown cell lines. FACS analysis revealed the activated CD8+ T cell population (CD8+ IFN-γ+) was significantly increased in both the primary tumor and metastatic lung samples of knockdown mice compared to controls (Fig 5 D&E). This increase was not observed in the spleen (S8 Fig C), demonstrating that the effect was specific to the primary tumor and metastatic sites. CD4+ Foxp3+ T regulatory cells were also affected by *Rnaseh2c* knockdown, but their reduction was restricted to the metastatic lung at the time of tissue harvest; NK cell infiltration was not affected (S8 Fig A&B). Together these data show that the immune system plays an important role in *Rnaseh2c*-mediated changes in metastasis.

### IRF3 activation, the proposed mechanism of AGS, is not observed upon *Rnaseh2c* knockdown in Mvt1 cells

Given that AGS also has an inflammatory component, we tested whether the current mechanism of this disease could also apply to our metastasis model. Recent AGS research has suggested that the cytosolic DNA-sensing cGAS-STING pathway is activated via IFR3 signaling in patients and mouse models of AGS [39–41]. This pathway leads to activation of Type I interferons and the production of interferon-stimulated genes to generate an immune response. We measured the production of a panel of interferon-related genes – including a subset known to be increased in AGS patients – in our *Rnaseh2c* knockdown cells using qRT-PCR. Of the 10 genes that were tested, 3 interferon-related genes showed a trending (p < 0.10) or significant (p < 0.05) increase in one or both knockdown cell lines (Fig 6A). These data indicate that *Rnaseh2c* knockdown results in a subtle upregulation of certain interferon-stimulated genes *in vitro*. To determine whether IRF3 is inducing these expression changes, IRF3 activation was examined. Although IRF3 could be phosphorylated and translocate to the nucleus in response to treatment with the STING activator DMXAA, these effects were not observed in response to *Rnaseh2c* knockdown alone (Fig 6B). NF-κB activation, which can also be induced by cGAS-STING signaling, was also examined. However, neither the canonical nor non-canonical NF-κB signaling pathways were shown to be activated (S9 Fig A&B).

**Fig 6.**
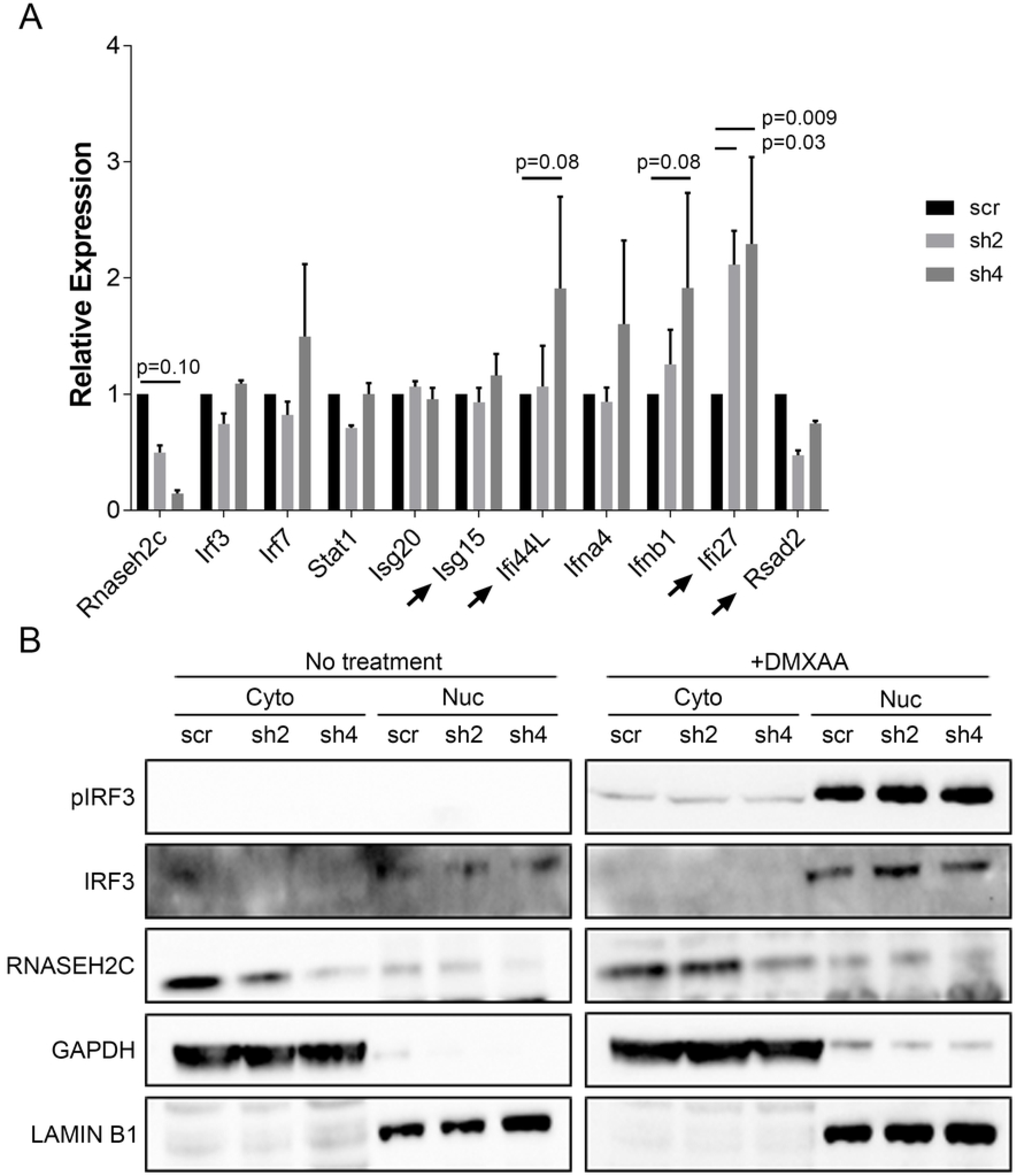
IRF3 signaling, the proposed AGS mechanism, is not activated in Mvt1 *Rnaseh2c* knockdown cells. (A) qPCR analysis of a panel of interferon stimulated genes in response to *Rnaseh2c* knockdown. Arrows indicate genes that are upregulated in AGS. Average ± standard error of three experiments. (B) Western blot analysis of IRF3 phosphorylation and nuclear translocation in *Rnaseh2c* knockdown cells following fractionation of the cytoplasmic (Cyto) and nuclear (Nuc) fractions. Sting activation using treatment with DMXAA was used as a positive control.

Double stranded RNA (dsRNA) can also activate an immune response by activating downstream signaling through IRF3 and IRF7 [42]. Since we had already determined IRF3 and NF-κB were not activated, we checked IRF7 as another potential transcription factor for dsRNA signaling. IRF7 was not observed to be enriched in the nucleus (S9 Fig C), suggesting that it is not actively signaling in response to *Rnaseh2c* knockdown. Therefore, further analysis of dsRNA was not pursued. These data suggest that the mechanism of immune activation in AGS may differ from that of the metastasis-associated *Rnaseh2c* effects.

### Immune response is not due to neoantigens or repetitive elements, suggesting a potentially enzyme-independent function

Because endogenous retroelements and noncoding RNAs are reported to be increased with dysfunctional RNase H2 and can act as nucleic acid triggers for immune activation [39, 43], we tested for changes in transcription of these molecules. For endogenous retroelements, qRT-PCR analysis of LINE-1 (L1) transcripts produced inconsistent results between shRNAs and between biological replicates (S9 Fig D), thus further analysis was not pursued. For noncoding RNAs, total RNA-sequencing of scramble control and sh4 knockdown cell lines was performed. Comparison of significantly differentially expressed noncoding RNAs produced a list of only 20 transcripts of three types: 14 antisense RNAs, 4 long intergenic noncoding RNAs, and 2 processed transcripts (S1 Table). Notably, this analysis also did not identify any differences in transcription of L1 elements. Since these results did not suggest a widespread dysregulation of noncoding RNA expression, the observed immune response is likely not due to recognition of these other RNA species.

Finally, it has been reported that RNase H2 mutations in both mouse fibroblasts and yeast can result in extensive DNA mutations and genomic damage [35, 44, 45]. Additionally, recent research has revealed the important role for cytotoxic T cells in response to neoantigens in tumors with high mutation burdens, such as melanoma [46]. We thus hypothesized that an increased burden of DNA mutations in *Rnaseh2c* knockdown cells might be generating tumor neoantigens targeted by cytotoxic T cells. To test whether the mutation burden was increased upon *Rnaseh2c* knockdown, we performed exome sequencing on metastases from nude mice. These samples were selected to enrich for cells capable of completing the metastatic cascade that might otherwise be cleared by the adaptive immune response in immunocompetent mice. Comparing exonic variant counts between control and knockdown metastases showed that there was not an increase in mutation burden with dysfunctional RNase H2 (S9 Fig E), suggesting neoantigens were likely not increased in the *Rnaseh2c* knockdown cells. While we cannot rule out that one of the few variants may be having an effect, it does not appear that there are extensive changes to these processes upon *Rnaseh2c* knockdown.

Together, these data eliminate numerous possible activators of an immune response based on known effects of RNase H2 dysfunction. Given these results, we propose that RNASEH2C may have a novel enzyme-independent activity that, when reduced in the context of breast cancer, results in reduced metastasis by engaging the immune system.

## Discussion

Metastasis is the primary cause of cancer-related deaths in breast cancer, yet the etiology of this lethal process is not well understood. In this study, we used a genetic strategy with the MMTV-PyMT model of spontaneous mammary tumorigenesis and metastasis to identify genes with differences in expression that associated with changes in metastasis. We demonstrated that *Rnaseh2c* functions as a metastasis modifier whose reduced expression activates a cytotoxic T cell-mediated immune response. These results further validate the meiotic genetics strategy we employ to identify candidate metastasis susceptibility genes. In addition, it demonstrates the ability of this approach to identify genes that impact metastasis through tumor non-autonomous mechanisms, such as interaction with the immune system.

Our data suggest that the *Rnaseh2c*-mediated metastatic effect is independent of its enzyme function since knockdown of *Rnaseh2c* affected metastasis, while knockdown of the catalytic subunit *Rnaseh2a* did not. This was surprising given that no other functions are known for the subunit. However, not all the AGS mutations identified in the RNase H2 genes affect enzymatic activity either. Although some mutations have been shown to reduce RNase H2 function - including the RNASEH2A G37S and RNASEH2C R69W mutations – other AGS mutations exhibit normal RNase H2 function [33, 44, 47–49]. This may indicate that a non-enzymatic function for the RNase H2 subunits, either in a complex or individually, could also be at play in AGS. The possibility of non-enzymatic functions for these subunits, especially in the context of cancer, has been suggested previously [50]. It should be noted, however, that the extent of the effect on enzyme activity was different between the gene knockdowns, and that a threshold of RNase H2 enzymatic activity may exist for the metastatic effects to be observed. A recent study using a ribonucleotide excision-deficient (RED) mutant of RNase H2 found that a single wildtype *Rnaseh2a* allele, and thus 50% enzyme activity, was sufficient to remove ribonucleotides from the genomic DNA and produce mice without any detectible abnormalities [51]. Interestingly, when RNase H2 activity was tested using the *Rnaseh2a*^+/RED^ embryo extracts, RER function was measured to be 43±6.6% of wildtype with hybrid activity like wildtype [51]. This is similar to the amount of residual RNase H2 RER activity measured in our *Rnaseh2c* knockdown cells. These data may suggest that the level of enzyme activity in *Rnaseh2c* knockdown cells is sufficient to remove incorporated ribonucleotides *in vivo* and, thus, the effects we observed on metastasis may not be due to the reduced enzyme activity.

Interestingly, although the results here implicate an *Rnaseh2c*-mediated immune response for metastatic progression, no convincing evidence of the AGS-associated cGAS-STING activation was observed upon *Rnaseh2c* knockdown. These observations suggest that the mechanism governing the immune activation in AGS may be different from the mechanism at play in breast cancer and metastasis, despite immune activation being common to the two diseases. In addition, the experiments that identified the involvement of cGAS-STING in AGS were done using mutations [39] or with knockout of a subunit [40] that were known to affect enzyme activity, thus it is not known whether this mechanism applies in a scenario of reduced expression or to enzyme-independent effects. It is also possible that the Interferon-stimulated gene activation is resulting from signaling of a non-canonical anti-viral response that was not tested here. Alternatively, these data may suggest that our *in vitro* condition does not fully recapitulate the *in vivo* environment of the mouse tumor or metastatic site. There could be other crucial molecules or cell types needed to amplify the signal generated by the knockdown, without which the signal is below the level of detection *in vitro*. Addressing this will require development of novel experimental techniques to study these biological effects *in vivo* as the cells undergo tumor progression and metastasis.

Interestingly, neither AGS patients nor carriers of the AGS mutations exhibit an observed increase in cancer incidence [52], though the early mortality of patients may help to explain this observation. However, multiple studies have reported associations between the RNase H2 genes and different types of cancer. *RNASEH2A* was identified as a susceptibility gene for aggressive prostate cancer [20] and, in another report, data from Oncomine revealed increased expression of RNASEH2A in a set of human cancers compared to normal tissue [18]. Similarly, analysis of overexpressed genes in gastric adenocarcinoma revealed increased expression of *RNASEH2B* [19]. In colorectal cancer, *RNASEH2C* mRNA was found to be reduced in tumor tissue compared to normal, while increased expression of *RNASEH2B* and *RNASEH2C* was correlated with metastasis [21], like in our study. Similar to what was observed here, no correlation between *RNASEH2C* expression and type I interferon (specifically *IFNB1*), which is typically increased in AGS patients, was observed [21]. A recent report also concluded that RNase H2 functions as a tumor suppressor in colorectal cancer using a mouse model containing H2B/p53 deletion in intestinal epithelial cells [23]. *Rnaseh2b* inactivation in the epidermis also resulted in increased proliferation, DNA damage, and development of squamous cell carcinoma [24]. Interestingly, this was accompanied by leukocyte infiltration which was shown to be independent of concomitant type I IFN production. Finally, in the context of breast cancer, a study analyzing low-frequency coding variants in loci identified by a genome-wide association study reported a significant association between a single amino acid change in RNASEH2C and breast cancer risk [22]. These data implicate the RNase H2 genes not only in the context of breast cancer, but also in progression of other cancer types. They also suggest that the effects of RNase H2 dysfunction and varied RNase H2 gene expression may be tissue-specific. Notably, many of the *in vivo* studies have used a knockout of the H2B subunit; while this is known to reduce the expression/stability of the H2A and H2C subunits, it may also leave residual H2C protein free to enact a tumor-promoting function, either in concert with or independent from the effects of absent H2B protein. Understanding how the biology of RNase H2 in cancer differs from its function in non-cancerous cells and how that biology differs across tissues may help shed light on additional targets for therapy.

The results from our study reveal an important role for the immune system in limiting *Rnaseh2c*-mediated metastatic disease through the cytotoxic T cell response. The involvement of the immune system may also help explain the results of the experimental (tail vein) metastasis assay. Unlike in spontaneous metastasis assays where the immune system is exposed to the developing tumor and thus primed to attack metastasizing cells, the immune system of the mouse receiving cells by tail vein injection has not been exposed to these cells prior to their bolus introduction into circulation and arrival in the lung. Therefore, the tumor-induced priming of the pulmonary immune microenvironment seen by spontaneously metastasizing cells would be absent in the experimental metastasis setting. In addition, the immune response at the primary tumor may be preventing cells from escaping, thus also ultimately preventing lung colonization. Both interpretations fit the observed results of the metastasis assays; additional study will be needed to determine whether either is accurate.

The involvement of the cytotoxic immune response introduces the possibility for effective immunotherapy in patients with reduced *RNASEH2C* expression, specifically immune checkpoint therapy. Immune checkpoint blockade would facilitate the continued targeting of the metastatic cells and primary tumors by naturally recruited cytotoxic T cells. This approach is currently being investigated in breast cancer (reviewed in [53]), but additional biomarkers such as *RNASEH2C* may help oncologists determine which patients are most likely to respond.

In summary, the results from our study implicate *Rnaseh2c* as a metastasis susceptibility gene that influences breast cancer metastasis through a novel, potentially enzyme-independent mechanism that stimulates cytotoxic T cells. This interaction supports the use of immune checkpoint inhibitors in the treatment of certain breast cancers. The classification of this gene as a metastasis modifier also adds to the list of known metastasis susceptibility genes that have the potential to be used by oncologists to identify patients with increased risk for developing metastatic disease, the primary cause of mortality in breast cancer.

## Materials and Methods

### Mouse strain analysis

The Diversity Outcross x MMTV-PyMT cross has been described previously [29]. Briefly, 50 female Diversity Outcross mice from Generation 5 were purchased from Jackson Laboratories and bred to male MMTV-PyMT mice. One hundred and fifteen transgene-positive female animals were generated and aged for mammary tumor and metastasis induction. When animals approached humane endpoints, they were euthanized by cervical dislocation after anesthetization by Avertin administration at 50 μL/g of body weight. Tumor tissue was collected, and surface pulmonary metastases enumerated as previously described [7]. Genotyping was performed with the MUGA mouse genotyping chip using spleen DNA. Genotypes for the 8 DO progenitor strains was downloaded from http://csbio.unc.edu/CCstatus/index.py and used to calculate linkage disequilibrium (LD) blocks for the DO animals using the Haploview software package. Haplotypes for each animal were then created using the PHASE software package (version 2.1) based on the Haploview LD blocks. Association analysis between the genotyping and the phenotypes was performed by using glm function in R. Associations were considered significant if the FDR < 0.05. For tumor gene expression, RNA was isolated from non-necrotic tumor samples using Trizol (Invitrogen) according to the manufacturer’s instructions. RNA was subsequently arrayed on Affymetrix Mouse Gene 1.0 × ST chip microarrays by the NCI Laboratory of Molecular Technology. The gene expression data is available through the Gene Expression Omnibus, accession no. GSE48566 [29]. Identification of genes within the haplotypes of interest having significant correlations with metastasis in the DO cross was performed using BRB-ArrayTools Version: 4.3.0 - Beta_1. Gene lists for each haplotype were used to filter the expression data, which were subsequently tested for correlation with metastasis using the BRB-ArrayTools Survival function, permuting the data 10000 times. Associations between gene expression and metastasis were considered significant if the permutated p-value < 0.05.

This study utilized the high-performance computational capabilities of the Biowulf Linux cluster at the National Institutes of Health, Bethesda, MD (http://biowulf.nih.gov).

### Cell lines and cell culturing

Mouse mammary carcinoma cell lines used were Mvt1 and 4T1. Cells were cultured in full DMEM (10% fetal bovine serum (FBS), 1% Penicillin and Streptomycin (P/S), 1% glutamine) unless otherwise described, and antibiotics for selection were included as indicated. Cells were incubated at 37°C with 5% CO_2_.

### Plasmid constructs

#### shRNA knockdown constructs

TRC lentiviral shRNA constructs against *Rnaseh2c* and *Rnaseh2a* were obtained from Open Biosystems (now Dharmacon) as glycerol stocks. The sequences for the shRNAs were as follows:

*Rnaseh2c* (Accession: NM_026616.2)
sh1: TRCN0000190101 - AAACCCAGGGCTGCCTTGGAAAAG
sh2: TRCN0000192140 - AAACCCAGGGCTGCCTTGGAAAAG
sh3: TRCN0000189664 - AAACCCAGGGCTGCCTTGGAAAAG
sh4: TRCN0000189847 - AAACCCAGGGCTGCCTTGGAAAAG
*Rnaseh2a* (Accession: NM_027187)
sh1: TRCN0000119583 - TTTGGTCTTGGGATCATTGGG
sh2: TRCN0000119582 - AACTGAACCGTACAAACTGGG
sh3: TRCN0000119585 - AAAGTGCTGTTGTAATCGAGC
sh4: TRCN0000119586 - AAACTGCCAGTTCTTCACAGC
sh5: TRCN0000119584 - AACTCAGACGACAACGACCCG

#### Overexpression constructs

MGC Mouse *Rnaseh2c* cDNA was purchased from Open Biosystems (now Dharmacon) as a glycerol stock (Accession: BC024333 Clone ID: 5040687). *Rnaseh2c* cDNA was amplified using Phusion polymerase and directional TOPO-cloning primers Fwd-5’-CACCATGAAGAACCCGGAGGAAG-3’ and Rev-5’-TCAGTCCTCAGGGACCT-3’. DNA was purified using the QiaQuick Gel extraction kit (Qiagen) and inserted into a Gateway entry vector using LR clonase (Invitrogen) according to the manufacturer’s protocol with an overnight ligation reaction.

### Virus transduction

1 × 10^6^ HEK293T cells were plated in 6 cm dishes 24 hr prior to transfection in P/S-free 10% FBS DMEM media. Cells were transfected with 1 ug of shRNA/cDNA and 1 ug of viral packaging plasmids (250ng pMD2.G and 750ng psPAX2) using 6 ul of Xtreme Gene 9 transfection reagent (Roche). After 24 hr, media was refreshed with full DMEM. The following day, virus-containing supernatant was passed through a 45 μm filter to obtain viral particles, which were then transferred to 100,000 target cells plated previously. 24 hr post-transduction the viral media was replaced with fresh full DMEM. Finally, 48 hr after transduction, the cells were selected with full DMEM containing 10 ug/ml puromycin (knockdown) or 5 ug/ml blasticidin (overexpression) and maintained in culture under selection.

### *In vivo* metastasis experiments

This study was performed under the Animal Study Protocol LCBG-004 approved by the NCI Bethesda Animal Care and Use Committee. Female virgin FVB/nJ or BALB/cJ mice were obtained from Jackson Laboratory at ~6 weeks of age. Athymic NCI nu/nu mice were obtained from NCI Frederick at the same age. Two days prior to injection, cells were plated at 1 million cells/condition into T-75 flasks (Corning) in non-selective DMEM media. A total of 100,000 cells per mouse were injected into the fourth mammary fat pad of the appropriate mouse strain. Mice were euthanized between 28 and 31 days post-injection by cervical dislocation following anesthesia by Avertin. Primary tumors were resected and weighed, and pulmonary surface metastases counted. For experimental metastasis assays, 100,000 tumor cells were injected into the tail vein of FVB mice. At 29 days post-injection, the mice were euthanized and pulmonary surface metastases counted. All surface metastasis counts were performed by a single investigator. Statistical analysis was performed using GraphPad Prism Software (La Jolla, CA) and R.

### Cell proliferation, cell cycle progression, and MTT assay

5000 cells/well were plated in 24-well plates (Corning, Inc.) with 6 technical replicates each. Cells were incubated in the IncuCyte Kinetic Live Cell Imaging System (Essen BioScience) at 37°C with 5% CO2 for up to 164 hr with imaging every 3-4 hr of 4 fields per well. Samples were imaged until either growth plateaued or cells reached 100% confluence, whichever occurred first. Data analysis was conducted using IncuCyte 2011A software. Representative experiment is shown of 4 biological replicates.

For cell cycle progression, cells were plated at equal numbers one day prior to collection. Cells were labeled with 10 μM BrdU for 25 min, then collected and 4.5 mL of ethanol was added in a dropwise manner while vortexing half-maximally. Cells were incubated for 30 min at 4 °C, then pelleted to remove supernatant. Cells were treated with 200 μL RNase A for 30 min, then in 500 μL of 5M HCl with 0.5% Triton X-100 for 20 min. Samples were neutralized in 10 mL of 1M Tris, washed in PBST, and incubated in 100uL of FITC-conjugated anti-BrdU (eBiosciences) solution (1:40 in 1% BSA in PBST) for 30 min mixing every 10 min. Cells were washed twice in PBST and resuspended in PBS with 20 ug/mL of propidium iodide. Samples were analyzed using FACS and FloJo software.

For sensitivity to doxorubicin treatment, cells were plated in a 96-well plate and grown overnight to ~70% confluency. The next day, cells were treated with increasing concentrations of doxorubicin for 24 hr. 50 ug per well of MTT solution in PBS was added to the media and incubated for 1 hr. Following incubation, MTT solution was removed and 100 μL MTT solvent dissolved in DMSO was added per well. The plate was placed on an orbital shaker with gentle agitation for 10 min protected from light, then absorbance read at 570 nm.

### Immunofluorescence

Cells were grown to approximately 50% confluency on 12 mm diameter cover slips and incubated overnight. Cells were washed twice in PBS and fixed with 4% formaldehyde or ice-cold methanol for 15 min, followed by two washes. Cells were treated with 10 mM glycine for 20 min and washed twice. Cells were then permeabilized with 0.1% Triton-X for 20 min, washed at least 3 times, and blocked with 5% goat serum for 1 hr followed by a wash in 0.1% BSA in PBS. Antibodies were prepared by dilution in 0.1% BSA in PBS and spun at 4 °C for 10 min at maximum speed. For staining, cells were incubated with primary antibody for 1 hr at room temperature to overnight at 4 °C followed by a wash in dilution buffer, followed by incubation in secondary antibody for 1 hr at room temperature. Cells were washed in dilution buffer, counter stained with DAPI, and mounted on slides for visualization. Antibodies used were as follows: yH2AX (1:100, Cell Signaling, #9718), S9.6 (1:200, Millipore, MABE1095).

### RNA isolation, reverse transcription, and qRT-PCR

Cells were plated at equal numbers one day prior to collection. RNA was isolated using TriPure (Roche) and reverse transcribed using iScript (Bio-Rad). Real-Time PCR was performed using VeriQuest SYBR Green qPCR Master Mix (Affymetrix). Expression of mRNA was defined by the threshold cycle and normalized to expression of Peptidylprolyl isomerase B (Ppib). Relative expression was calculated using the delta Ct method. Primer sequences are provided in the supplementary information (S2 Table).

### Western blotting

Cells were plated at equal numbers one day prior to collection. Protein was extracted on ice from collected cells using Golden Lysis Buffer (10 mM Tris pH 8.0, 400 mM NaCl, 1% Triton X-100, 10% Glycerol plus oComplete protease inhibitor cocktail (Roche) and phosphatase inhibitor (Sigma)). Protein concentration was measured using Pierce’s BCA Protein Assay Kit and analyzed on the Versamax spectrophotometer at a wavelength of 560 nm. Equal amounts of protein were combined with NuPage LDS Sample Buffer and NuPage Reducing Agent (Invitrogen) and were run on NuPage Bis-Tris gels in MOPS buffer. Proteins were transferred onto a PVDF membrane (Millipore), blocked in 5% milk in 0.05% TBST (TBS + 5% Tween) for 1 hr and incubated in the primary antibody (in 5% milk) overnight at 4 °C. Membranes were washed with TBST and secondary antibody incubations were done at room temperature for 1 hr. Proteins were visualized using Amersham ECL Prime Western Blotting Detection Reagent (GE Healthcare) using film or the ChemiDoc Touch Imaging System (BioRad). Densitometry analysis was performed using ImageJ or Image Lab Software (BioRad). Antibodies used were as follows: RNASEH2C (1:1000; Abcam #ab89726), RNASEH2A (1:1000; Proteintech #16132-1-AP), BETA-ACTIN (1:10,000; Abcam #ab6276), anti-FLAG (1:1000; Sigma #080M6034), RNASEH1 (1:1000; Proteintech #15606-1-AP), caspase 3 (1:1000; Cell Signaling #9662), cleaved caspase 3 (1:1000; Cell Signaling #9661), p21 (1:333; C-19; Santa Cruz #sc-397), pIRF3 (1:1000; Cell Signaling #29047), IRF3 (1:1000; Millipore Sigma #MABF268), GAPDH (1:10,000; Millipore #MAB374), LAMIN B1 (1:1000; Abcam #ab16048-100), IRF7 (1:1000, Abcam #ab109255), p65/RELA (1:1000; Cell Signaling #8242), IκBα (1:1000; Cell Signaling #4814), NF-κB1 p105/p50 (1:1000; Cell Signaling #12540), NF-κB2 p100/p52 (1:1000; Cell Signaling #4882), RELB (1:1000; Cell Signaling #10544), Histone H3 (1:10,000; Abcam #ab1791), goat anti-rabbit (1:10,000; Santa Cruz), anti-rabbit (1:3000; Cell Signaling #7074), goat anti-mouse (1:10,000; GE Healthcare NA931).

### *In vitro* RNase H2 activity assay

RNase H2 activity was determined as described in [51]. Briefly, 0.1 mg/mL of protein lysate from Mvt1 cells with manipulated *Rnaseh2c* or *Rnaseh2a* expression were combined with 1 μM of either a 5’-^32^P-labeled 12 base DNA_5_-RNA_1_-DNA_6_ hybridized with complementary DNA for RER activity or a uniformly labeled poly-^32^P-rA/poly-dT substrate for hybrid activity. Reaction products were separated using a 12 or 20% TBE-urea polyacrylamide gel and quantified using a Typhoon imager.

### Immunophenotyping

Tumors, metastatic lungs, and spleens were collected from FVB mice (n=5 per group) 30 days after mammary fat pad injection. Tumors and lungs were dissociated in 5mL PBS using the MACS dissociator and passed through a 100 μM filter. Spleens were mashed through a 0.40 μM filter, treated with ACK lysis buffer on ice for 5 min, and resuspended in RPMI. Lymphocytes were isolated from tumor and lung tissue using Lympholyte-M Cell Separation Media (Cedar Lane). Briefly, pelleted tissue digest was resuspended with RPMI followed by the addition of Lympholyte-M. Samples were centrifuged and supernatant spun again to isolate lymphocytes which were resuspended in RPMI. Lymphocytes were activated with Leukocyte Activation Cocktail with Golgi Plug (BD Pharmigen) for 4 hr at 37 °C. Following activation, immune cells were stained with a fixable live/dead stain (Invitrogen) in PBS followed by surface antibody staining for 20 min in FACS buffer (PBS with 0.5% BSA and 0.1% sodium azide). For intracellular cytokine staining, cells were first stained for surface markers (Thy1.1; CD4; CD8; NK1.1) and later stained for intracellular molecules (IFN-γ; Foxp3) following fixation and permeabilization according to the manufacturer’s protocols (BD Cytofix/Cytoperm). Data were collected on the LSRII- Fortessa (BD) and analyzed with FlowJo software (Tree Star). Antibodies used were as follows at 1:200 dilution: Thy1.1 (Ebioscience, #25-0900-82), CD4 (BD, #553729), CD8 (Biolegend, #100750), Interferon gamma (BD, #557998), FoxP3 (Ebioscience, #17-5773-82), NK1.1 (Ebioscience, 12-597182).

### Sequencing and analysis

RNA was extracted from small sections of primary tumor tissue using TriPure, followed by organic extraction with chloroform and precipitation by isopropanol. RNA was then purified using the RNeasy^®^ Mini Kit (Qiagen) with on-column DNase digestion. RNA quality was tested using the Agilent 2200 TapeStation electrophoresis system, and samples with an RNA integrity number (RIN) score >7 were sent to the Sequencing Facility at Frederick National Laboratory. Preparation of mRNA libraries and mRNA sequencing was performed by the Sequencing Facility using the HiSeq2500 instrument with Illumina TruSeq v4 chemistry. Sample reads were trimmed to remove adapters and low-quality bases using Trimmomatic software and aligned with the reference mouse mm9 genome and Ensemble v70 transcripts using Tophat software. RNA mapping statistics were calculated using Picard software. Library complexity was measured by unique fragments in the mapped reads using Picard’s MarkDuplicate utility. Analysis of mRNA-seq data was performed using the Partek Genomics Suite. GSEA was performed using the annotated genes ordered by log_2_ fold change with 1000 permutations. Gene sets with a nominal p-value of < 0.05 and a false discovery rate (FDR) of < 0.25 were considered significantly enriched. Gene ontology pathway analysis was performed using Ingenuity Pathway Analysis (Qiagen) with genes that exhibited at least two-fold change in expression in either direction. Both analyses were performed comparing sh4 to scramble tumors.

In silico analysis of immune cell infiltration by mRNA-seq data was performed using ImmQuant software [38]. Briefly, a list of differentially expressed genes generated from mRNA-sequencing analysis was used along with the software-provided reference dataset (ImmGen consortia, [54]) and marker genes for mouse. Relative expression was calculated using the scramble tumors designated as the control sample.

Exome sequencing was performed using DNA isolated from spontaneous metastases in NCI nu/nu mice. Four metastases were isolated from lungs of mice 30 days after orthotopic tumor implantation as described above. Metastases were dissociated using a sterile disposable pestle in a microcentrifuge tube in tail lysis buffer (100 mM Tris pH8.5, 5 mM EDTA, 0.2% SDS, 200 mM NaCl). 160 ug of Proteinase K (Invitrogen) was added to the tube and contents were incubated at 55 °C overnight then for an additional 1hr with shaking at 1400 rpm. Genomic DNA was then isolated from the dissociated metastases using the DNA miniprep kit (Zymo Research) according to the manufacturer’s instructions. DNA quality was tested using the Agilent 2200 TapeStation electrophoresis system, and samples were sent to the Sequencing Facility at Frederick National Laboratory. Variant analysis was performed on the NIH High Performance Computer system, Helix & Biowulf. Sequencing reads were mapped using BWA and/or Bowtie using the mouse mm10 reference genome. Variant calling was performed using mpileup and/or GATK HaplotypeCaller. Germline variants in the Mouse Genome Project (mgp.v5) were filtered out; low quality variants (quality score ≤ 30) were also removed. Remaining variants were annotated using Annovar to identify nonsynonymous single nucleotide variants and indels. In-house R scripts were used for statistical analyses.

RNA for total RNA sequencing was isolated as described above and sent to the Sequencing Facility at Frederick National Laboratory. To analyze expression of long noncoding RNAs, sequence data was aligned to the Long non-coding RNA gene annotation from GENCODE (version M15, available at https://www.gencodegenes.org/mouse/) using Tophat software. DESeq2 was used to compare control and knockdown samples.

### Immunohistochemistry

Fresh tumor tissue was fixed in 10% phosphate buffered formalin for 24 hr then transferred to 70% ethanol. Paraffin embedding and sectioning was performed by Histoserv, Inc. (Germantown, MD). Tumor sections were rehydrated by sequential 5 min incubations in Histochoice (Sigma), 100% ethanol, 90% ethanol, 70% ethanol, and water. Slides were then incubated in 3% H_2_O_2_ for 5 min. Antigen retrieval was performed using Antigen Retrieval Citra (Biogenex) and heating for 10 min, followed by blocking for 25 min with Serum-free Protein Block (Dako). Primary antibody incubation was performed at room temperature for 1 hr in a humidified chamber. Antibodies used were Ki67 (1:200; Abcam), cleaved caspase 3 (1:250; Cell Signaling). After washing, incubation with ImmPRESS™ HRP Anti-Rabbit IgG (Peroxidase) Polymer Detection Kit (Vector) was performed for 30 min at room temperature in a humidified chamber. DAB chromogen solution was applied for 4.5 min, followed by counterstain for 3 min. Slides were dehydrated by sequential incubations in the reverse order reported above for rehydration. Coverslips were affixed, and slides were scanned using the Aperio Slide Scanning System (Leica Biosystems). Positive nuclei (Ki67) and stained area (cleaved caspase 3) were analyzed using the ImageScope software (Leica Biosystems). Quantification is average of 3 sections, 1 from each of 3 independent tumors.

### Statistical Analysis

Metastasis assays requiring multiple comparisons was analyzed using the Kruskal-Wallis test (non-parametric one-way Anova) with post-hoc Conover-Inman correction for multiple analyses. The Z-method was used to combine p-values with weighting as appropriate. RNase H2 assays were analyzed using a one-tailed T test. Ki67 and cleaved caspase 3 IHC were analyzed using the Kruskal-Wallis test. Cell cycle progression was analyzed using a one-way Anova of knockdown sample compared to scramble. Immunophenotyping was analyzed using the Mann-Whitney test. ISG expression was analyzed using a two-way Anova with Dunnett’s multiple comparison correction. Statistical significance was evaluated as a p-value < 0.05.

## Acknowledgments

The authors would like to thank present and past members of the Hunter Laboratory for technical assistance and critical review of the manuscript. We would also like to thank the members of Laboratory of Cancer Biology and Genetics at the NCI for helpful discussions and shared reagents. We thank Helen Michael for IHC assistance, Aleksandra Michalowski for assistance with statistical analysis, and the NCI Animal Facility staff. We thank Steve Shema and Qin Wei in the CCR Genomics Core and Langston Lim in the CCR Confocal Microscopy Core for their expertise. We thank Bao Tran and Jyoti Shetty at the Sequencing Facility at Frederick National Laboratory for their expertise. This research was supported in part by the Intramural Research Program of the NIH at the National Cancer Institute, Center for Cancer Research and National Institute of Child Health and Human Development.

## Supporting Information

**S1 Fig. A haplotype on mouse chromosome 19 predicts distant metastasis-free survival in HER2 enriched patients.**

(A) Metastasis-associated haplotypes identified using the DO × PyMT cross described in Fig 1A. Plus sign denotes the haplotype selected for further investigation. (B) 5-gene signature of the chromosome 19 haplotype selected in (A) and their weights calculated using the mouse hazard ratios. (C) Kaplan-Meier analysis of distant metastasis-free survival (DMFS) of patients from the GOBO dataset stratified into terciles of the cumulative signature score of the gene signature in (B). Scores are calculated in GOBO by considering both the direction of expression and relative contribution of the gene to the signature as represented by the signature weight. HR-hazard ratio

**S2 Fig. qRT-PCR analysis of shRNA-mediated gene knockdowns in the Mvt1 cell line.**

(A) qRT-PCR analysis of *Rnaseh2c* expression following shRNA-mediated knockdown in Mvt1 cells. Average ± standard error of three experiments. (B) qRT-PCR analysis of *Rnaseh2c* expression following transduction of Mvt1 cells with an exogenous expression construct. Average ± standard deviation of three experiments. (C) qRT-PCR analysis of *Rnaseh2a* expression following shRNA-mediated knockdown in Mvt1 cells. Average ± standard deviation of three experiments.

**S3 Fig. Knockdown of *Rnaseh2c* reduces pulmonary metastasis in the 4T1 cell line.**

(A) qRT-PCR analysis of *Rnaseh2c* expression following shRNA-mediated knockdown in 4T1 cells. (B) RNASEH2C protein expression by western blot. One representative experiment is shown. (C-E) Spontaneous metastasis of 4T1 knockdown lines sh2 and sh4 was assessed as described. Tumor mass (C) and pulmonary metastases (D) were quantified at euthanasia and normalized (metastases per gram of tumor, E); average ± standard deviation, n = 10 mice per group.

**S4 Fig. *Rnaseh2c* knockdown does not affect proliferation, apoptosis, or sensitivity to doxorubicin.**

(A) *In vitro* cellular confluence was monitored as an indirect measurement of proliferation using the IncuCyte imaging system; average ± standard deviation of six technical replicates. (B) Full length and cleaved caspase 3 analysis in *Rnaseh2c* knockdown cells by western blot. (C) Ki67 (top) and cleaved caspase 3 (bottom) staining by IHC of tumor sections, representative image of staining three independent tumors. Quantification is shown in Fig 4A. (*D*) *Rnaseh2c* knockdown cells were treated with increasing concentrations of doxorubicin over 24 hours and cell viability was measured using the MTT assay. Absorbance at 570nm is reported as a percentage of the untreated condition.

**S5 Fig. *Rnaseh2c* knockdown does not produce double-strand DNA breaks.**

Immunofluorescence staining of γ-H2AX in Mvt1 cells with *Rnaseh2c* knockdown. Cells were grown to approximately 50% confluency on glass coverslips for staining. One of two independent experiments is shown. Magnification, 63X.

**S6 Fig. *Rnaseh1* expression compensates for *Rnaseh2c* knockdown.**

(A) Immunofluorescence staining of RNA/DNA hybrids using the S9.6 antibody in Mvt1 cells with *Rnaseh2c* knockdown. One of three independent experiments is shown. Magnification 100X. (B) RNASEH1 protein expression upon *Rnaseh2c* knockdown. Densitometry relative to Actin for three independent experiments is reported below. (C) Percent RNA/DNA hybrid (RNase H) activity in Mvt1 cells with knockdown of *Rnaseh2c*. Activity levels are presented relative to untransduced Mvt1 cells. Average ± standard deviation of two technical replicates. NS – not significant

**S7 Fig. *In silico* analysis of immune cell-specific gene expression patterns predicts infiltration of knockdown tumors by CD8+ T cells.**

mRNA-sequencing data was analyzed using ImmQuant software for changes in immune cell-specific gene expression and compared to reference gene expression profiles from defined inflammatory states. Predicted presence of immune cell types identified in the sh4 tumors are reported between −1 (dark blue, lowest presence) and 1 (dark red, highest presence) compared to scramble control tumors. Categories of immune cells are shown in yellow.

**S8 Fig. CD4+ T regulatory cells and NK cells do not exhibit the same pattern as CD8+ cytotoxic T cells.**

Immunophenotyping of cells within the primary tumor (left) or metastatic lungs (right) at euthanasia: (A) Average percent T regulatory cells identified by CD4+ Foxp3+ staining. (B) Average percent natural killer (NK) cells identified by NK1.1 staining. (C) Presence of activated (IFN-γ producing) CD8+ T cells in the spleen at euthanasia. Average ± SEM; NS – not significant

**S9 Fig. Other known immune-related pathways are not activated in *Rnaseh2c* knockdown cells.**

(A) Western blot analysis of canonical NF-κB signaling using fractionated (top) and whole cell (bottom) lysate from *Rnaseh2c* knockdown cells. (B) Western blot analysis of noncanonical NF-κB signaling using fractionated (top) and whole cell (bottom) lysate from *Rnaseh2c* knockdown cells. (C) Western blot analysis of IRF7 nuclear translocation in the *Rnaseh2c* knockdown cells following fractionation of the cytoplasmic (Cyto) and nuclear (Nuc) fractions. (D) Analysis of three families of L1 elements by qRT-PCR. Average ± standard deviation of three experiments. (E) Exome sequencing to analyze mutation burden (number of sequence variants) following *Rnaseh2c* knockdown. Average ± standard deviation; n = 4 metastases per group.

**S1 File. Full IPA analysis from Mvt1 *Rnaseh2c* sh4 versus scramble control RNA-sequencing**

**S1 Table. Differentially expressed non-coding RNAs in Mvt1 *Rnaseh2c* sh4 knockdown versus scramble control cells by total RNA-sequencing**

**S2 Table. Primers for qRT-PCR and cloning**

